# Oomycete effector AVRblb2 targets cyclic nucleotide-gated channels through calcium sensors to suppress pattern-triggered immunity

**DOI:** 10.1101/2023.01.18.524664

**Authors:** Soeui Lee, Hye-Young Lee, Hui Jeong Kang, Ye-Eun Seo, Joo Hyun Lee, Doil Choi

## Abstract

Transient, rapid increase of cytosolic Ca^2+^ upon pathogen infection is essential for plant pathogen-associated molecular pattern (PAMP)-triggered immunity (PTI). Several cyclic nucleotide-gated channels (CNGCs) have been implicated; however, their regulatory mechanisms remain elusive. Here, the *Phytophthora infestans* RXLR effector AVRblb2 family targeted NbCNGC18–20 at the plasma membrane, inhibiting Ca^2+^ influx and PTI. AVRblb2 required calmodulin (CaM) and calmodulin-like (CML) proteins as co-factors to interact with *N. benthamiana* CNGCs (NbCNGCs), forming the AVRblb2-CaM/CML-NbCNGCs complex. After recognizing PAMPs, NbCNGC18 formed active heteromeric channels with other CNGCs, potentially providing selectivity for diverse signals to fine-tune cytosolic Ca^2+^ levels and responses. AVRblb2 suppressed the Ca^2+^ influx and oxidative burst induced by NbCNGC18 heteromeric complexes. Silencing CNGC18, CNGC20, and CNGC25 compromised the effect of AVRblb2 on *P. infestans* virulence, confirming that AVRblb2 contributed to virulence by targeting CNGCs. Our findings delineated the regulatory mechanism and role of effector-targeted Ca^2+^ channels in plant innate immunity.

## INTRODUCTION

The plant immune system is a sophisticated, multilayered system that protects against pathogens (Boller and Felix, 2009; Jones and Dangl, 2006). When pathogens invade, the plant immune system is activated through the recognition of conserved microbial molecules known as pathogen-associated molecular patterns (PAMPs) by the pattern-recognition receptor, resulting in the activation of PAMP-triggered immunity (PTI). However, pathogens have evolved to secrete effector proteins that can disrupt PTI responses, leading to effector-triggered susceptibility. To counteract this disruption, the plant immune system employs intracellular nucleotide-binding leucine-rich-repeat (NLR) receptors that recognize its cognate pathogen effectors and induce effector-triggered immunity (ETI) (Cui et al., 2015; Jones and Dangl, 2006; Kadota et al., 2019). Both PTI and ETI trigger a range of downstream responses, including calcium influx, reactive oxygen and nitrogen species production, activation of mitogen-activated protein kinase (MAPK) cascades, transcriptional reprogramming, and hormone signaling (Cui et al., 2015). PTI and ETI can also potentiate each other to provide full resistance to pathogen invasion (Yuan et al., 2021)

An increase in cytosolic calcium concentrations is a critical early immune signaling event (Grant et al., 2000; Moeder et al., 2019). Ca^2+^ is a universal second messenger in eukaryotes, and its cytosolic levels are typically maintained at low levels (∼10^−8^ to 10^−7^ M) by sequestration in intracellular stores, such as vacuoles, the endoplasmic reticulum, and apoplasts, via active transport. Upon PAMP recognition, calcium channels at the plasma membrane are activated, leading to transient Ca^2+^ influx decoded by a variety of Ca^2+^ sensors containing EF-hand motifs, including the calmodulin (CaM) and calmodulin-like (CML) proteins. Increased Ca^2+^ influx activates respiratory burst oxidase homolog, an NADPH oxidase that generates a burst of reactive oxygen species (ROS) and triggers downstream immune responses (Xu et al., 2022).

Cyclic nucleotide-gated channels (CNGCs) are well-studied calcium channels implicated in plant immune responses. These channels have a cytosolic C-terminus with a CaM binding domain, a cyclic nucleotide-binding domain, and several phosphorylation sites. Recent studies discovered several regulatory mechanisms of CNGCs via the cytosolic C-terminus (DeFalco et al., 2016; Fischer et al., 2017; Tian et al., 2019). In Arabidopsis, the extensively studied AtCNGC2 and AtCNGC4 contribute to plant immunity by increasing cytosolic Ca^2+^ (Moeder et al., 2019; Wang et al., 2019). CNGC2 and CNGC4 form a heteromeric complex kept inactive by CaM binding; however, the resting state can be released by the receptor-like cytoplasmic kinase, Botrytis-induced kinase 1, during PAMP-induced Ca^2+^ bursts (Tian et al., 2019). Furthermore, AtCNGC11/CNGC12 and AtCNGC19/20 have been linked to immunity via an autoimmune mutant that mediates constitutive Ca^2+^ influx (Yoshioka et al., 2006; Yu et al., 2019). However, the precise molecular mechanisms by which pathogens disrupt Ca^2+^ signaling to sustain their life cycles during early PTI are largely unknown.

Plant pathogens have evolved a plethora of effectors to disrupt host defense signaling pathways and enhance their invasion. *Phytophthora infestans*, an oomycetes pathogen, secretes hundreds of effector proteins, including RXLR family members (Whisson et al., 2007; Zheng et al., 2014). Among these, the AVRblb2 gene family is particularly intriguing because it is found in all *P. infestans* isolates and is conserved in other pathogenic *Phytophthora spp*., playing a critical role in pathogen virulence (Oliva et al., 2015). AVRblb2 and its homologs target various host proteins, including the cysteine protease C14, a potato MAPK cascade protein (StMKK1), and CaM (Bozkurt et al., 2011; Du et al., 2021; Naveed et al., 2019). Furthermore, most of the multiple copies of AVRblb2 are specifically recognized by Rpi-blb2 (Oh et al., 2009), showing the importance of this gene family in the co-evolution of *P. infestans* and their host plants. Similar to AVRblb2, several pathogen effectors directly target the main Ca^2+^ sensor protein, CaM. For example, the *P. infestans* RXLR effector Suppressor of early flg22-induced Immune response 5 (SFI5) targets SlCaM, and the SFI5-CaM interaction is necessary for effector function (Zheng et al., 2018). Furthermore, the *Pseudomonas syringae* effector HopE1 targets the microtubule-associated protein 65 (MAP65) in a CaM-dependent manner to inhibit pathogenesis-related protein 1 (PR-1) protein secretion (Guo et al., 2016). However, the role of effectors in the host immune response in the context of calcium signaling remains unknown.

In this study, we showed that the *P. infestans* RXLR effector AVRblb2 family could suppress PTI by targeting NbCNGCs at the plasma membrane. AVRblb2 targeted a subset of NbCNGCs (CNGC18–20) via the Ca^2+^ sensors CaM and CML36, forming the AVRblb2-CaM/CML36-CNGC complex. Molecular studies demonstrated that AVRblb2 inhibited flg22-induced Ca^2+^ influx by preventing the dissociation of CaM/CML36 and NbCNGC18. Following flg22 treatment, CNGC18 formed a heteromeric complex with other CNGCs that might be targeted by AVRblb2. AVRblb2 function was disrupted in the silencing of CNGC3, CNGC11, CNGC18, and CNGC20 plants, demonstrating that the function of AVRblb2 in virulence required its association with multiple CNGCs. Our findings have provided a working model for heteromeric CNGC complexes in the fine-tuning of Ca^2+^ homeostasis in plant immunity and identified how pathogen effectors disturb PTI.

## RESULTS

### AVRblb2 interacts with CaM and CML36

The calmodulin-binding domain (CBD, aa 78–82 in the C-terminal region of AVRblb2) of AVRblb2 is required for the AVRblb2-CaM interaction (Naveed *et al*., 2019). We used the CBD deletion (AVRblb2(S)) and alanine substitution (AVRblb2(A)) mutants described by Naveed et al. (Naveed *et al*., 2019) to investigate the effect of CaM binding to AVRblb2 on pathogenicity. When *P. infestans* was inoculated onto *N. benthamiana* leaves expressing each construct, 3FLAG-AVRblb2-expressing leaves showed increased susceptibility to infection compared to the GST control leaves, as previously reported (Bozkurt *et al*., 2011). However, the 3FLAG-AVRblb2(A)- and 3FLAG-AVRblb2(S)-expressing leaves had compromised susceptibility compared to 3FLAG-AVRblb2, indicating that CaM binding is required for AVRblb2 to function as an effector (Fig. 1A, B).

**Figure 1.**
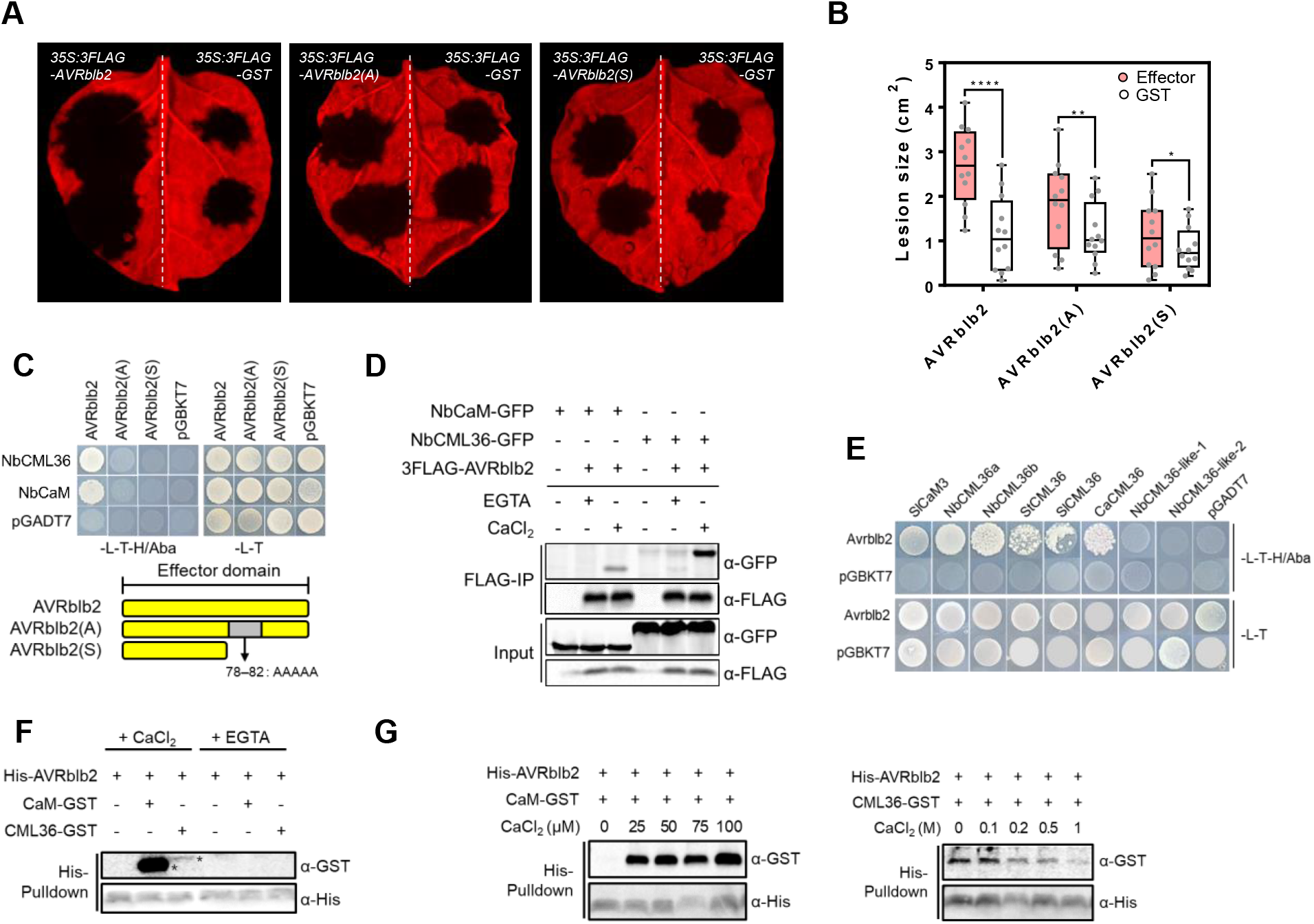
AVRblb2 interacts with NbCaM and NbCML36. (A and B) *P. infestans* T30-4-infected *N. benthamiana* leaves expressing GST, AVRblb2, AVRblb2(S) and AVRblb2(A) 6 days after infection (A). Lesion size (B) of inoculation sites shown in (A). Values are mean ± SD of the three biological replicates (n=18). Significance was determined using a t-test. ***, P < 0.001; **, P<0.001; *, P <0.05. (C) Growth of yeast cells co-expressing GAL4 BD-AVRblb2 with GAL4 AD-NbCaM or -NbCML36 on synthetic media lacking LTH or LT (upper panel). Schematic representation of the AVRblb2 mutants used in (A and B) (lower panel). (D) AVRblb2 *in planta* associates with NbCaM or NbCML36 dependent on calcium ion. Total proteins extracted from *N. benthamiana* leaves expressing 3FLAG:AVRblb2 with NbCaM-GFP or NbCML36-GFP were subjected to FLAG IP using α-FLAG agarose in the presence of 1 mM of CaCl_2_ or 2 mM EGTA. AVRblb2-CaM/CML36 were detected by western blot using anti-GFP antibodies. (E) Growth of yeast cells co-expressing GAL4 BD-AVRblb2 with GAL4 AD-CML36 homologs of *N. benthamiana*, tomato and pepper on synthetic media lacking LTH or LT. (F and G) *In vitro* pull-down assay of His-AVRblb2 using His-affinity resin. Crude bacterial extracts from *E. coli* expressing His-AVRblb2 with CaM2-GST or CML36-GST were subjected to pull-down assays with 1 mM CaCl_2_ or 1 mM EGTA (F). Pull-down assay between His-AVRblb2 and CaM-GST or CML36-GST with serial concentration of CaCl_2_.

In addition to CaM, *Solanaceae* plants have many calmodulin-like proteins (CMLs) in their genomes that play important roles as Ca^2+^ sensors. In the *Nicotiana benthamiana* genome, there are 55 annotated CML genes with structures similar to canonical CaM (Zhao et al., 2013). Using cDNA library screening, we found that *N. benthamiana* NbCML36a/b interacted with AVRblb2. Moreover, both CaM and CML36a interacted with AVRblb2 in the yeast two-hybrid (Y2H) system. However, AVRblb2(A) and AVRblb2(S) did not, indicating that the CBD of AVRblb2 is required for its interactions with CaM and CML36 (Fig. 1C). These interactions were confirmed by co-immunoprecipitation (co-IP) (Fig. 1D). Because CaM and CML36 are calcium sensors, we added calcium or Ca^2+^ chelator EGTA to the total protein extracts used for IP to determine how calcium affected the interactions between AVRblb2 and CaM or CML36. The results showed that AVRblb2 interacted with CaM and CML only when calcium was present. To see if CML36 orthologs in *Solanaceae* plants also interacted with AVRblb2, CaCML36, SlCML36, and StCML36 were tested using the Y2H assay (Fig. 1E). All tested CML36s interacted with AVRblb2, whereas NbCML36-like-1 and NbCML36-like-2 did not, indicating that AVRblb2 specifically interacts with CML36 orthologues. To better understand the interactions of CaM and CML36 with AVRblb2, we used His:AVRblb2, CaM-GST, and CML36-GST in an *in vitro* competition assay. CaM-GST interacted strongly with His:AVRblb2 in the presence of calcium, whereas CML36-GST had only a weak interaction (Fig. 1F). In particular, the CaM binding affinity for AVRblb2 increased as the calcium concentration increased, whereas CML36 interacted with AVRblb2 only in the low Ca^2+^concentrations (Fig. 1G), indicating that AVRblb2 binding to CaM and CML36 had different affinity depending on Ca^2+^ concentration, leading to different roles for AVRblb2 based on these calcium sensors.

### AVRblb2 associates with NbCNGC18 in a Ca2+ sensor-dependent manner

Because CaM and CML36 bind to calcium channels and regulate calcium signaling through these channels, we hypothesized that AVRblb2 might associate with calcium channels via CaM and CML36 to regulate calcium signaling. Thus, we performed co-IP assays with AVRblb2 and *N. benthamiana* CNGC family members. We cloned 9 CNGCs from the *N. benthamiana* genome (Saand et al., 2015) (NbCNGC3, 4, 5, 7, 18, 19, 20, 21 and 25) to determine which ones interacted with AVRblb2 (Fig. S1). We found that NbCNGC18, NbCNGC19, and NbCNGC20 specifically interacted with AVRblb2 when co-expressed in *N. benthamiana*. (Fig. 2A). NbCNGC18 and NbCNGC19 share over 98% amino acid sequence identity, with identical C-terminal cytosolic regions. Both NbCNGC18 and NbCNGC19 showed strong interactions with AVRblb2.

**Figure 2.**
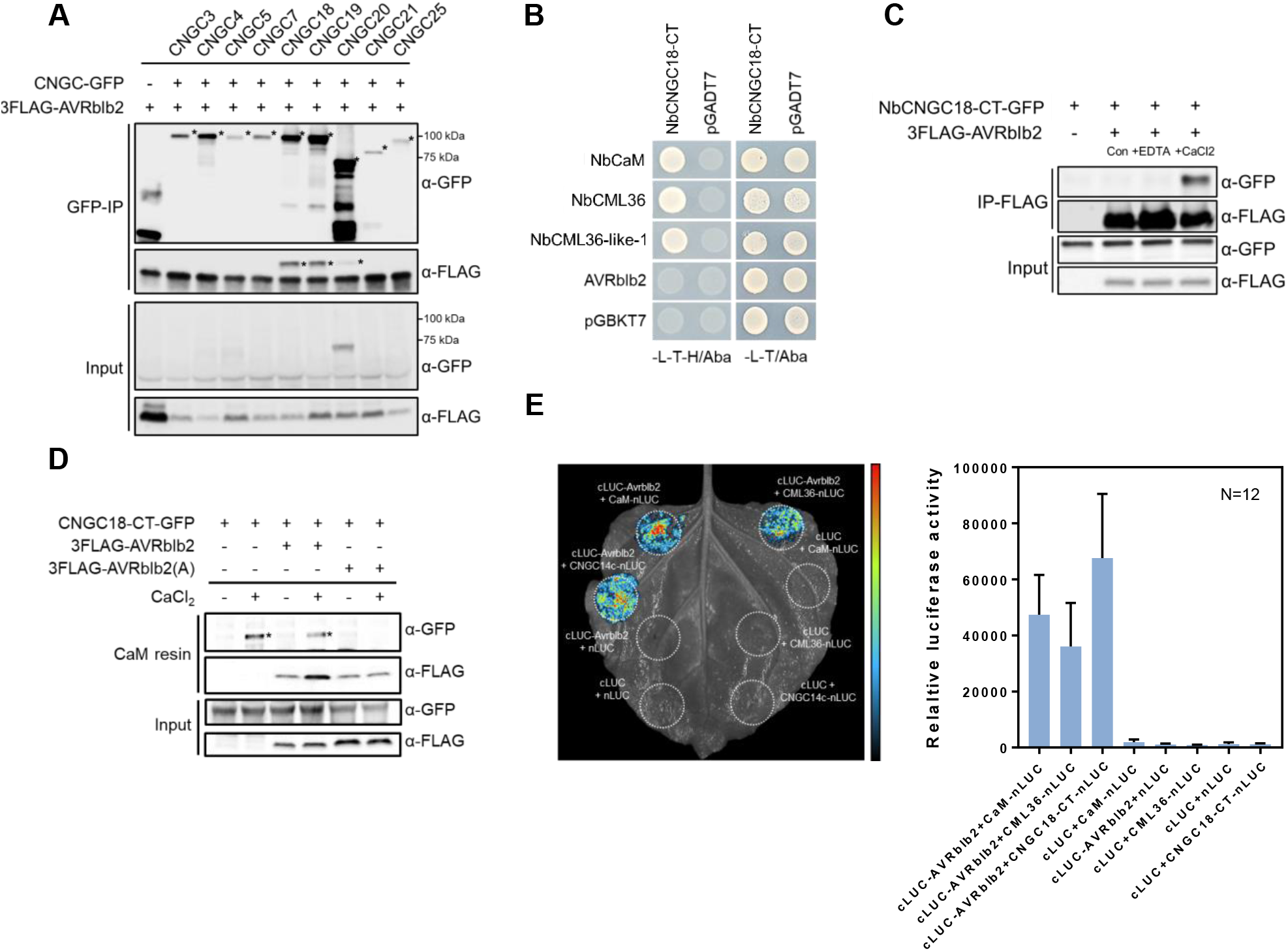
AVRblb2 associates with NbCNGCs through calcium sensors. (A) 3FLAG-AVRblb2 and NbCNGCs-GFP were transiently expressed in *N. benthamiana*, and protein extracts were subjected to co-IP using anti GFP agarose. Precipitated proteins were detected by western blotting using anti FLAG antibody. (B) The interaction of CNGC18-CT with NbCaM, NbCML36s or AVRblb2 in a yeast two-hybrid assay. Yeast cells were plated onto SD-LT or SD-LTH with aureobasidin A (Aba) and pictured 3 days after dropping. Empty pGBKT7 and pGADT7 were used as a negative control. (C) *In planta* co-IP assay on *N. benthamiana* expressing AVRblb2 and NbCNGC18-CT with 1 mM EGTA or 1 mM CaCl_2_. Con represents a mock-treated sample. (D) AVRblb2 interacts with the NbCNGC18-CT in an *in planta* pull-down assay. Total proteins extracted from *N. benthamiana* leaves expressing CNGC18-CT-GFP and 3FLAG-AVRblb2 or 3FLAG-AVRblb2(A) were incubated with CaM. The resins were pelleted for western blotting with anti-GFP or anti-FLAG antibody respectively. (E) Split-luciferase complementation assay to determine interaction between AVRblb2 and NbCML36 or NbCNGC18-CT in *N. benthamiana*. The photos were taken at 2 day post infiltration (right panel) and luminescence was quantified by the emitted light units (left panel). Values are mean ± SD (n=12). NbCaM-nLuc with cLuc-AVRblb2 was used as a positive control.

Because of the high identity between these two effectors, NbCNGC18 was chosen for further study. CaM, CML36, and CML36-like-1, interacted with NbCNGC18 (NbCNGC18-CT) in the Y2H system, whereas AVRblb2 did not (Fig. 2B), implying that AVRblb2 interacted indirectly with NbCNGC. Co-IP with additional CaCl_2_ or EGTA was performed to confirm the interaction between AVRblb2 and CNGC18-CT *in planta*. The AVRblb2 and NbCNGC18-CT interaction was only detected in the presence of Ca^2+^, indicating that this interaction was Ca^2+^-dependent (Fig. 2C). The combination of AVRblb2 and NbCNGC18-CT restored the catalytic activity of luciferase in *N. benthamiana* in the split-luciferase assay, confirming that NbCNGC18 associates with AVRblb2 *in planta* (Fig. 2E). We also investigated whether AVRblb2 forms a calcium-dependent complex with CaM/CML36 and NbCNGC18-CT. To test this possibility, we used CaM affinity resin to bind proteins with EF-hand motifs *in planta* using pull-down assays. CaM resin precipitated both AVRblb2 and CNGC18-CT (Fig. 2D), suggesting that NbCNGC18 interacted with AVRblb2 via CaM/CML36. These findings show that the association between AVRblb2 and NbCNGC18 occurs via CaM/CML36 in a Ca^2+^-dependent manner. Furthermore, these results indicated that AVRblb2 might influence CNGC activity via CaM and CML36.

### AVRblb2 suppresses PTI-induced Ca2+ influx

Several CNGCs contribute to the transient Ca^2+^ influx induced by PTI and ETI (Zhao et al., 2021). Thus, we hypothesized that AVRblb2 could regulate Ca^2+^ signaling induced by PAMP through NbCNGC binding. We tracked Ca^2+^ influx in response to flg22 treatment by monitoring cytosolic Ca^2+^ dynamics in *N. benthamiana* plants expressing the fluorescent protein-based Ca^2+^ sensor YC3.6. Cytoplasmic Ca^2+^ levels peaked at 15 to 20 min after flg22 treatment and then gradually declined in the GST-expressing conditions. AVRblb2 strongly inhibited flg22-induced Ca^2+^ influx, whereas AVRblb2(A) only weakly inhibited this influx (Fig. 3A-1). AVRblb2 family also suppressed Ca^2+^ influx (Fig. S2). Furthermore, AVRblb2 broadly suppressed elevated cytosolic Ca^2+^ levels induced by elf18 and chitin (Fig. S3), indicating that AVRblb2 inhibited PAMP-induced Ca^2+^ influx in *N. benthamiana*, disrupting the PTI response.

**Figure 3.**
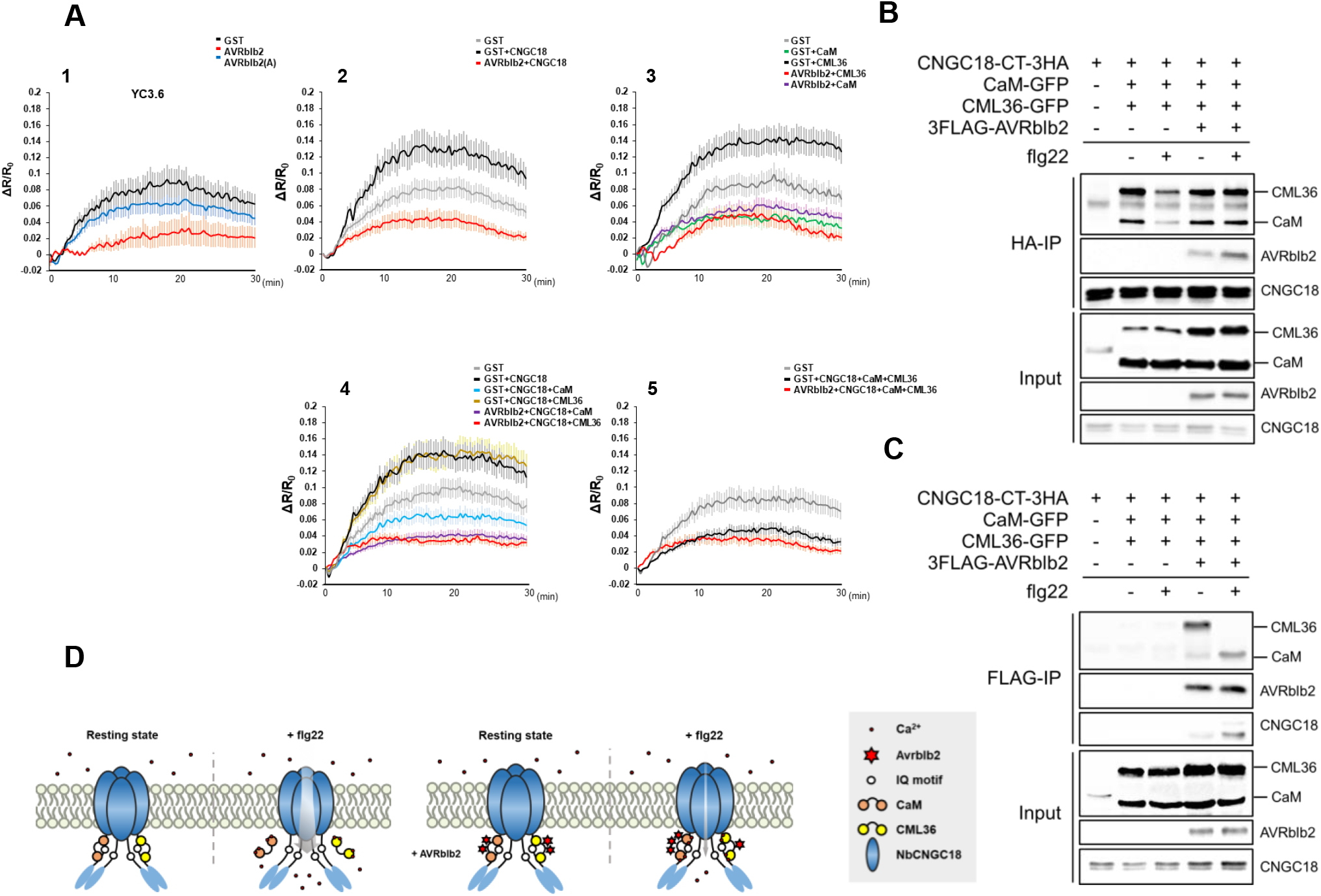
AVRblb2 regulates NbCNGC18 activity via calcium sensors. (A) Calcium sensor-CNGC association affects flg22-mediated Ca2+ influx in *N. benthamiana*. Dynamics of [Ca2+]cyt in the YC3.6 transgenic *N. benthamiana* leaves were monitored in time course. FRET ratio represents [Ca2+]cyt in leaves transiently expressing indicated protein(s) in response to 1 μM flg22 at time = 0 s. (n ≥ 20). (B) Co-IP experiments confirm that suppressed interaction between calcium sensors and CNGC18-CT in response to flg22 was restored in presence of AVRblb2. Combinations of GFP-tagged CaM and CML36, FLAG-tagged AVRblb2 and HA-tagged CNGC18-CT were transiently expressed in *N. benthamiana* leaves by agroinfiltration, as indicated. (C) CML36 and CaM were co-IPed with AVRblb2 and CaM-AVRblb2 association is enhanced upon elicitor treatment. (D) Working model for calcium sensors and CNGC-mediated Ca2+ influx in response to PAMP. In the resting state, NbCNGC18 associates with CaM and CML36. Upon perception of PAMP, CaM and CML36 are separated from NbCNGC18, which results in activation of NbCNGC18 to induce Ca2+ influx. When PTI is suppressed in presence of AVRblb2, disassociation of CaM/CML36-NbCNGC18 is inhibited by formation of complex AVRblb2-calcium sensors-CNGC18.

We also examined Ca^2+^ influx under NbCNGC18-expressing conditions to see if NbCNGC18 was involved in PAMP-triggered calcium signaling. When NbNCGC18 was overexpressed, we saw a significant increase in Ca^2+^ influx after flg22 treatment but an impaired response in NbCNGC18-silenced plants (Fig. 3A-2, Fig. S4), indicating that NbCNGC18 positively regulated flg22-induced Ca^2+^ influx in *N. benthamiana*. The flg22-induced Ca^2+^ spike was strongly inhibited in CaM-expressing *N. benthamiana* leaves, whereas Ca^2+^ influx was significantly enhanced in CML36-expressing plants (Fig. 3A-3), demonstrating that CaM and CML36 had opposing roles in flg22-induced Ca^2+^ influx. To investigate the effect of CaM and CML36 on NbCNGC18 activity, we measured Ca^2+^ influx in plants co-expressing NbCNGC18 and CaM or CML36. CaM significantly reduced Ca^2+^ influx compared to CNGC18 alone, whereas leaves co-expressing CML36 and NbCNGC18 had similar Ca^2+^ influx levels as those expressing NbCNGC18 alone (Fig. 3A-4). Furthermore, co-expression of CaM, CML36, and NbCNGC18 showed lower Ca^2+^ peaks than the GST control, indicating that CaM-mediated suppression of the NbCNGC18-induced Ca^2+^ spike was not rescued by CML36 (Fig. 3A-5). Surprisingly, AVRblb2 expression inhibited flg22-induced Ca2+ influx in all the cases (Fig. 3A). These findings showed that NbCNGC18 was involved in the flg22-induced transient Ca^2+^ influx regulated by CaM and CML36, which had opposing functions. Moreover, AVRblb2 suppressed Ca^2+^ influx to disrupt the PTI response.

### AVRblb2 prevents the dissociation of CaM/CML36 from NbCNGC18

CaM and CML36 have different binding affinities for AVRblb2 (Fig. 1F) and have opposite effects on NbCNGC18 activity. Thus, we hypothesized that AVRblb2 could suppress NbCNGC18 activity by regulating the binding affinity of CaM and CML36 for NbCNGC18. To test this hypothesis, we performed co-IP experiments to examine the binding affinity between NbCNGC18 and CaM or CML36 in response to flg22 treatment. CaM and CML36 were both associated with NbCNGC18 in the resting state; however, flg22 treatment significantly reduced their binding to NbCNGC18. When AVRblb2 was present, CaM and CML36 could not be released from NbCNGC18 after flg22 treatment (Fig. 3B). This finding suggested that AVRblb2 might regulate NbCNGC18 activity by preventing the dissociation of CaM or CML36 and NbCNGC18. The CaM and CML36 affinities to AVRblb2 differed in response to flg22. AVRblb2 mostly interacted with CML36 in the absence of flg22 but bound to CaM after flg22 treatment (Fig. 3C), which was consistent with the *in vitro* pull-down data for AVRblb2 and CaM or CML36 in the presence of calcium (Fig. 1G) and indicated that AVRblb2 primarily associated with CaM in the presence of high Ca^2+^levels *in vitro* and *in planta* but interacted with CML36 under low Ca^2+^ conditions (Fig. 3D). These findings demonstrated that changes in the dynamics of the interactions between CaM, CML36, and NbCNGC18 were early events in PAMP perception. They also suggested that AVRblb2 primarily used CaM to suppress NbCNGC18 activity and disrupt the PTI response.

One of the major regulatory mechanisms for CNGC activity is phosphorylation at the C-terminus (Curran et al., 2011; Zhou et al., 2014). Recent studies revealed that upon pathogen attack, some CNGCs are activated following phosphorylation by receptor kinases or receptor-like kinases (Ladwig et al., 2015; Tian et al., 2019; Wang et al., 2019; Yu et al., 2019). We examined the levels of NbCNGC18 phosphorylation using p-Thr and p-Ser antibodies, which detect phosphorylated threonine and serine, respectively. We also used Phos-tag, which detects overall phosphorylation, to see if AVRblb2 regulated CNGC18 activity by mediating its phosphorylation. NbCNGC18-CT phosphorylation, particularly on Thr and Ser, increased in flg22-treated leaves (Fig. S5A–C). There was no difference in Thr and Ser phosphorylation in the presence of AVRblb2; In addition, the overall amount of phosphorylation in NbCNGC18-CT was similar to those in absence of AVRblb2, suggesting that AVRblb2 did not affect the phosphorylation status of NbCNGC18.

### AVRblb2 suppresses the activity of other CNGCs by interacting with CNGC18

Plant CNGCs form homomeric and heteromeric complexes under specific conditions, and each complex can affect channel activity (Brost et al., 2019; Chin et al., 2013; Dietrich et al., 2020; Pan et al., 2019; Tian et al., 2019; Tian et al., 2020). We used co-IP and liquid chromatography-tandem mass spectrometry (LC-MS/MS) analysis of NbCNGC18-expressing leaves after flg22 treatment to investigate the CNGC candidates interacting with NbCNGC18 during flg22-induced PTI. We identified NbCNGC3, NbCNGC11, NbCNGC20, and NbCNGC25 as putative NbCNGC18 interactors. These interactions were validated by co-IP in response to flg22 treatment. NbCNGC3, NbCNGC20, and NbCNGC25 interacted strongly with NbCNGC18 in response to flg22 treatment, whereas NbCNGC11 binding to NbCNGC18 remained unchanged (Fig. 4A). Furthermore, the interaction of NbCNGC18 to itself was reduced in response to flg22 (Fig. 4A), indicating that NbCNGC18 functions in a complex with other CNGCs in the presence of flg22 rather than itself.

**Figure 4.**
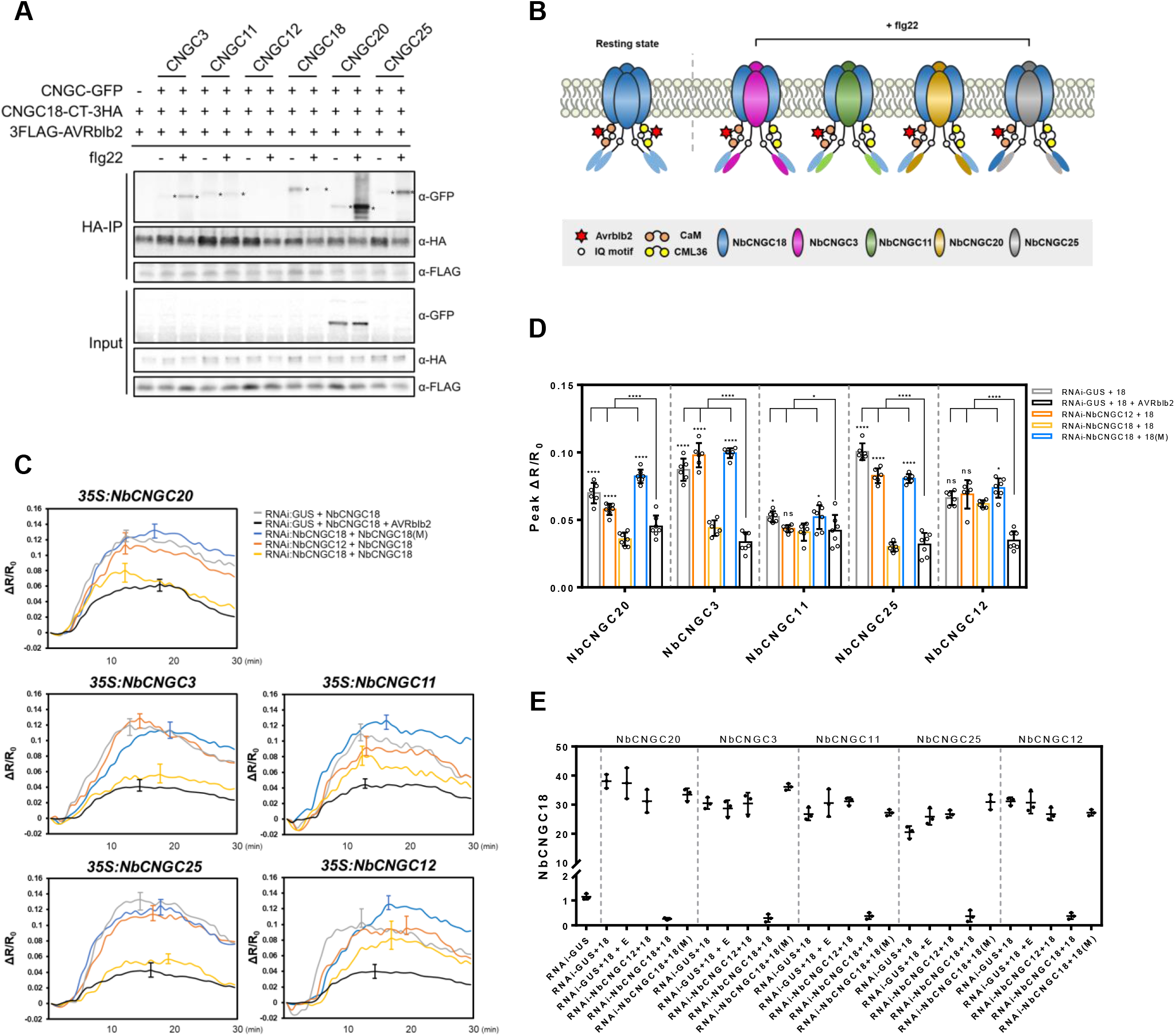
NbCNGC18 forms a heterotetrameric complex with other CNGCs during the PTI response. (A) NbCNGC18 interacts with CNGC3/20/25 after flg22 treatment. NbCNGC18-CT-3HA, NbCNGCs-GFP and 3FLAG-AVRblb2 were co-expressed in *N. benthamiana* by Agroinfiltration. Collected leave samples treated with H2O or 1 μm flg22 for 10 min were subjected to co-IP experiments. (B) Schematic model of NbCNGC18 heteromeric channels induced by flg22. (C) Co-overexpression of NbCNGC18 with NbCNGC3/20/25, but not NbCNGC11/12, enhanced Ca2+ influx upon flg22 treatment. (D) Peak value of cysotolic Ca2+ influx in leaf discs used in (C). Means ± standard deviation of the peak value are given from 18 replicates. (two-way ANOVA; Tukey’s HSD; P < 0.05). (E) Quantitative RT-PCR analysis of *NbCNGC18* gene expression. NbActin was used as an internal standard. Results shown are mean ± SE from triplicate technical replicates from one of three representative experiments with similar results. P-values were calculated using t-tests (*p < 0.005).

Bimolecular fluorescence complementation (BiFC) was used to confirm the interactions of NbCNGC18 with other CNGCs (Fig. S6A). BiFC was also used to investigate whether these interacting CNGCs could also interact with AVRblb2. In addition to NbCNGC19 and NbCNGC20, which belong to the same clade as NbCNGC18 (Fig. S3), NbCNGC3 and NbCNGC4 interacted with AVRblb2 (Fig. S6B). Notably, NbCNGC11 and NbCNGC25 did not interact with AVRblb2, despite showing a fluorescence signal in BiFC with NbCNGC18 (Fig. S6A, B). These findings demonstrated that NbCNGC18 could form heteromeric complexes with other CNGCs in response to PAMP perception and suggested that AVRblb2 mediated the formation of specific heteromeric CNGC complexes to suppress Ca^2+-^influx in early PTI (Fig. 4B).

Next, we investigated which heteromeric NbCNGC complex was involved in the flg22-induced PTI response with NbCNGC18. We generated a hairpin RNA interference (RNAi) construct against NbCNGC18 under the control of the CaMV 35S promoter (35S:dsNbCNGC18). NbCNGC12-RNAi was generated as a negative control. A codon-shuffled NbCNGC18 (NbCNGC18(M)) was generated to avoid RNAi complementation. When NbCNGC18 was co-expressed with NbCNGC3, NbCNGC20, or NbCNGC25, flg22-induced Ca^2+^ influx was significantly increased. In contrast, Ca^2+^ influx did not increase when NbCNGC18 was silenced, even though the other CNGCs were overexpressed. Furthermore, NbCNGC11 or NbCNGC12 overexpression did not affect Ca^2+^ influx (Fig. 4C–E), consistent with the co-IP results (Fig. 4A). Together, these findings suggest a working model in which NbCNGC18 forms heteromeric complexes with other NbCNGCs during flg22-induced PTI.

### AVRblb2 suppresses the PTI response and enhances virulence by regulating CNGCs

Because increased Ca^2+^ influx is required to activate PTI downstream signaling, we investigated the role of AVRblb2 in the flg22-triggered immune response. First, we used oxidative burst assays to determine ROS levels. AVRblb2 significantly reduced ROS production, whereas AVRblb2(A) caused a partial loss in ROS production (Fig. 6A). AVRblb2, but not AVRblb(A), suppressed the expression of PTI marker genes, such as *NbICS1, NbPR1, NbAcre31*, and *NbWRKY22* (Heese et al., 2007) (Fig. 6B). Furthermore, immunodetection of flg22-dependent activation of the MAPKs, correlated with ROS production, indicating that AVRblb2 suppressed the overall PTI response (Fig. 5A–C). Co-expression of AVRblb2 and NbCNGC18 also suppressed ROS production. Expression of NbCNGC18, co-expression of NbCNGC18 and NbCNGC20, and co-expression of NbCNGC18, NbCNGC20, and NbCNGC25 increased ROS production additively, whereas AVRblb2 disrupted the increase in ROS production caused by the overexpression of NbCNGCs (Fig. 5E).

**Figure 5.**
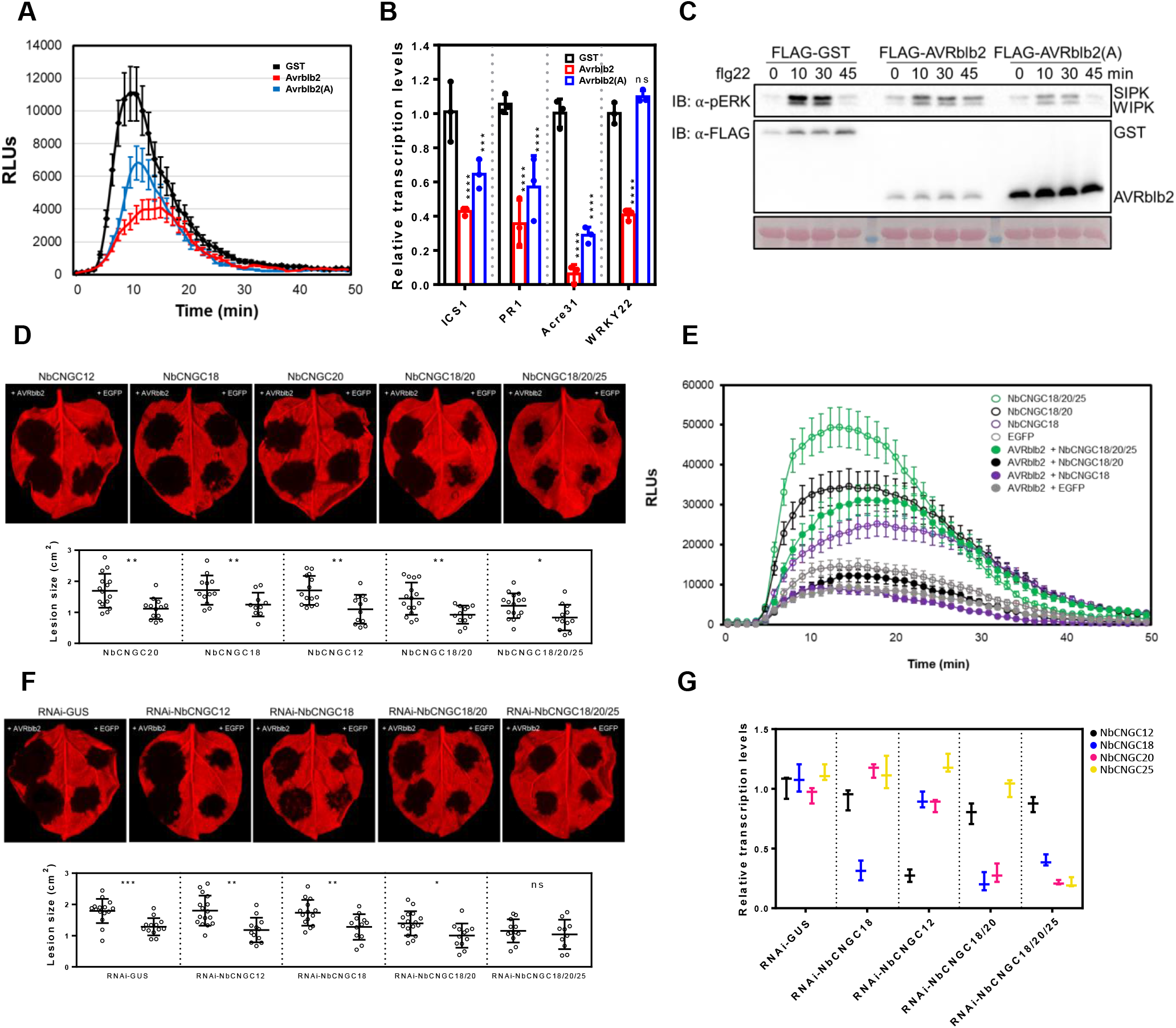
AVRblb2 suppresses the PTI response and enhances virulence by regulating the activity of NbCNGCs. (A) Flg22-induced ROS burst is strongly suppressed in AVRblb2 expressed leaves. ROS burst in *N. benthamiana* leaves after treatment with 1 μM flg22. Results shown are mean of 6 replicates ± SE (n=6). (B) Transcriptional induction of defense related genes by flg22 is suppressed in AVRblb2-expressed leaves. Gene expression of *NbICS1, NbPR1, NbAcre31* and *NbWRKY22* in *N. benthamaiana* upon flg22 treatment was analyzed by qRT-PCR. (C) Differential MPK activation in *N. benthamiana* expressing AVRblb2 or AVRblb2(A) after flg22 treatment. The kinetics of kinase activation is shown by immunoblot analysis using an anti-pERK antibody. (D) Leaves expressing AVRblb2 exhibit enhanced disease symptoms after *P. infestans*. The leaves expressing NbCNGCs and AVRblb2 were inoculated with droplets of zoospore suspension from P. infestans 2 days after the agroinfiltration (upper panel). EGFP was used as negative control for AVRblb2. Corresponding lesion sizes were shown in lower panel respectively. Values are mean ± SD of the three biological replicates (n=15). (E) Flg22-induced reactive oxygen species (ROS) burst in NbCNGCs-overexpressed plants. Leaf discs from 4-week-old *N. benthamiana* were treated with 1 μM flg22, followed by ROS measurement by chemi-luminescence assay. (F) Enhanced disease symptom induced by overexpression of AVRblb2 is compromised in CNGC18/20/25-silenced *N. benthamiana*. Lesion size (lower panel) of inoculation sites shown in upper panel. Corresponding lesion sizes were shown in lower panel respectively. Values are mean ± SD of the three biological replicates (n=15). (G) Quantitative RT-PCR analysis of NbCNGCs gene expression in RNAi-plants. *NbActin* was used as an internal standard. Results shown are mean ± SE from triplicate technical replicates.

Finally, we performed pathogen infection assays to investigate the effect of NbCNGCs on pathogenic resistance. Leaves co-expressing NbCNGC18, NbCNGC20, and NbCNGC25 had increased resistance to *P. infestans*, indicating Ca^2+^ influx caused by the overexpression of multiple NbCNGCs could enhance PTI (Fig. S7). But AVRblb2 expressed parts showed increased susceptibility when NbCNGC12, NbCNGC20 or multiple CNGCs were co-expressed than GST control; indicating that AVRblb2 could suppress the resistance induced by overexpression of NbCNGCs, which result is consistent with ROS assay (Fig. 5D). When we targeted NbCNGC18 with RNAi constructs (Fig. 6G), AVRblb2 retained its virulent activity, implying that NbCNGC18 was functionally redundant with NbCNGC19, NbCNGC20, or both NbCNGCs (Fig. 6F). However, AVRblb2 did not increase *P. infestans* susceptibility in leaves with silenced NbCNGC18, NbCNGC20, and NbCNGC25 (Fig. 6F). These results suggested that the presence of multiple CNGCs has an additive effect in boosting the immune defense against pathogens by forming heteromeric complexes with NbCNGC18. Together, AVRblb2 disrupts PTI response to enhance pathogen virulence through the regulation of NbCNGC18.

## DISCUSSION

Rapid elevation of cytosolic free Ca^2+^ levels in plant cells is an important early event in immune signaling. Previous research found that calcium sensors regulate CNGCs, which function as calcium channels for Ca^2+^ influx through the plasma membrane in response to PAMP (Tian *et al*., 2019; Yoshioka *et al*., 2006; Yu *et al*., 2019). However, it is unknown how pathogen effectors disrupt calcium signaling to suppress PTI. Here, we showed that the *P. infestans* effector AVRblb2 targeted NbCNGCs via the Ca^2+^ sensors CaM and CML36 to disrupt the PTI response by suppressing calcium influx, allowing the pathogen to colonize successfully. AVRblb2 and all tested AVRblb2 homologs interacted with NbCNGC18-CT in the presence of calcium and suppressed PAMP-induced calcium influx, indicating that they were functionally redundant with respect to *P. infestans* virulence.

CaM and CML36 likely have opposite roles in calcium channel regulation and calcium signaling in response to pathogens. CaM is released from NbCNGC18 in response to flg22 treatment, likely leading to the NbCNGC18 activation and increased calcium influx. However, in the presence of flg22, AVRblb2 increased the binding affinity of CaM for NbCNGC18 and disrupted their dissociation, resulting in suppressed calcium influx (Fig. S8). In contrast, CML36 increased calcium influx while interacting primarily with AVRblb2 under normal conditions. AVRblb2 family members are recognized by the broad-spectrum resistance protein Rpi-blb2, causing hypersensitive responses (HRs). We confirmed an enhanced Rpi-blb2-AVRblb2-mediated HR cell death when CML36 was co-expressed (data not shown), indicating that CML36 may monitor the interactions between AVRblb2 and CaM; upon recognition by Rpi-blb2, robust calcium influx leads to HR. However, how Rpi-blb2 recognizes AVRblb2 activity during calcium signaling needs further investigation.

Plant CNGCs form heteromeric complexes, and the combination of CNGCs varies depending on the signal. A previous study demonstrated heterotetrameric channel formation by AtCNGC2-AtCNGC4, AtCNGC7 or AtCNGC8-AtCNGC18 and AtCNGC19-AtCNGC20 *in planta*. However, it is unclear whether those CNGC-CNGC combinations are constitutive or dynamically regulated, depending on the context (Chin et al., 2013; Pan et al., 2019; Zhao et al., 2021). In this study, we discovered that NbCNGC18 existed as a homotetramer in the resting state but formed a heteromeric complex with specific CNGCs in response to PTI. Furthermore, AVRblb2 interacted with NbCNGC18 to suppress calcium channel activity, disrupting the PTI response. AVRblb2 still exhibited virulent activity under NbCNGC18-silenced conditions, implying that NbCNGC19 and NbCNGC20, which interact with AVRblb2, likely share functional redundancy with NbCNGC18. Indeed, AVRblb2 appeared to lose its virulence when NbCNGC18, NbCNGC20, and NbCNGC25 were silenced. However, we cannot rule out the possibility that other effectors work with other calcium channels or that other calcium channels are involved in the immune response. Given the expanded number of CNGC, CaM and CML gene family members in *N. benthamiana*, the vast array of CNGC combinations in tetrameric complexes with calcium sensors could explain the generation of unlimited number of stimulus-specific Ca^2+^ oscillations.

In summary, in the current study, we identified members of the defense-related Ca^2+^ channels as targets of a pathogen effector (AVRblb2) in *N. benthamiana* and developed a working mechanism for PAMP perception. Furthermore, we deciphered the role of this *P. infestans* effector in suppressing PAMP-induced Ca^2+^ influx during early PTI responses. Our findings could have far-reaching implications for future studies of Ca^2+^ signaling in plant immunity and understanding a pathogen’s strategy for manipulating early plant immune responses.

## Acknowledgments

We want to acknowledge Dr. E. Park (University of Wyoming, USA) for providing the RNAi construct. This work was supported by National Research Foundation of Korea (NRF) grants funded by the Korean government (MSIT) to D.C. (No. 2018R1A5A1023599 [SRC] and 2021R1A2B5B03001613). The authors have no conflicts of interest to declare.

## Author contributions

S.L., J.H.L., and D.C. wrote the paper and coordinated the research. S.L. performed experiments. J.H.L. performed the *in vitro* pull-down assay and assisted with the LC-MS/MS analyses. H.K. generated the *N. benthamiana* transgenic plants used in the study. H.Y.L. performed the split-luciferase assay, and Y.E.S. performed the MAPK assay.

## STAR methods

### Plant material and Agrobacterium-mediated transient overexpression

*N. benthamiana* plants were grown and maintained in a work-in chamber in a controlled environment with an average temperature of 23–24°C and long day conditions (16 hr light cycle). *A. tumefaciens* GV3101 was used to deliver T-DNA constructs into *N. benthamiana* leaves. Overnight cultures of *A. tumefaciens* were centrifuged at 3500 × g for 10 min, and suspensions were adjusted to a final OD600 of 0.3 in infiltration buffer (10 mM MgCl2, 10 mM MES, pH 5.6, and 150 μM acetosyringone).

### Plasmid construction

The AVRblb2 gene (PITG_20300) was cloned into the binary vector pCAMBIA2300-LIC using a ligation-independent cloning method (Oh et al., 2010). NbCaM, NbCML36, and all candidate NbCNGC genes were amplified from *N. benthamiana* cDNA and cloned into pCAMBIA2300-LIC containing a C-terminal GFP or HAx3 tag. The recombinant plasmids were introduced into *A. tumefaciens* strain GV3101.

### Ca2+ concentration measurements

The monitoring of changes in cytosolic calcium concentrations was performed using transgenic *N. benthamiana* plants expressing YC3.6 under the control of the cauliflower mosaic virus promoter 35S. Leaf disks from YC3.6 *N. benthamiana* transgenic line were transferred to a 96-well black plate and incubated in ddH2O at 25°C for 12 hr, followed by treatment with dH2O (Ctrl) or 1 μM flg22. FRET fluorescence was measured immediately upon treatment using a multi-label plate reader (Varioskan LUX, ThermoFisher, USA) with an excitation at 440/40 nm and emission detection at 520/40 nm. Measurements were recorded at 1 min intervals. The FRET-based cpVenus/ECFP ratio was used for the ratio (R) calculation (440/520) and normalized to the initial ratio (R0). The data were plotted versus time (ΔR/R0). Experiments were repeated ten times. One representative result is presented.

### Hairpin RNA-mediated gene silencing

The NbCNGC silencing fragment was amplified from *N. benthamiana* cDNA using the primers listed in Table S1. The specificity of the silencing fragment was analyzed using the *N. benthamiana* genome sequence and associated gene silencing target prediction tool (SGN VIGS tool: https://vigs.solgenomics.net). The purified amplicon was cloned into the pRNAi-GG vector as previously described (91). *N. benthamiana* leaves were infiltrated with Agrobacterium carrying pRNAi-GUS or various pRNAi-CNGC constructs at a final OD600 of 0.3.

### Protein extraction and immunoblotting

*N. benthamiana* leaves were ground in liquid nitrogen and mixed with protein extraction buffer (10% [v/v] glycerol, 25 mM Tris-HCl [pH 7.5], 1 mM EDTA, 150 mM NaCl, 1% [w/v] polyvinylpolypyrrolidone, and 1× protease inhibitor cocktail). Suspensions were mixed and centrifuged at 12000 × g for 15 min at 4°C. Total proteins were separated on 10% or 12% sodium dodecyl sulfate (SDS)-polyacrylamide gels and transferred to Immobilon-PSQ polyvinylidene difluoride membranes. The membranes were washed with PBST (PBS with 0.1% Tween 20) for 3 min and then blocked in 5% non-fat milk for 1 hr. The membranes were then incubated with rabbit monoclonal anti-GFP (1:12000, Invitrogen, A10260) or anti-FLAG (1:12000, Sigma-Aldrich, F3165) antibody for 2 h, followed by three washes with PBST. The membranes were then incubated with PBST for 10 min each before the addition of the secondary anti-mouse Ig-horseradish peroxidase (HRP) (1: 15,000, Abcam, ab6708) or anti-rabbit Ig-HRP (1:15,000, Abcam, ab6702) antibody for 1 h. After three washes with PBST, the blots were developed using enhanced chemiluminescence (Thermo Scientific, USA).

### Co-immunoprecipitation (co-IP)

Proteins were extracted from two *N. benthamiana* leaves at 1.5 days post-agroinfiltration by homogenizing the leaves in GTEN extraction buffer (10% [v/v] glycerol, 25 mM Tris-HCl [pH 7.5], 1 mM EDTA, 150 mM NaCl, 1% [w/v] polyvinylpolypyrrolidone, and 1× protease inhibitor cocktail) and centrifuging the homogenates at 12000 × g for 15 min. For co-IP, total protein extract (1 mg) was mixed with 10 μl GFP-Trap®-A agarose beads (Chromatek, Munich, Germany), anti-FLAG agarose beads (A7470, Sigma-Aldrich), or anti-HA affinity matrix beads (Roche) and incubated end-over-end for 2 hr at 4°C. Beads were washed five times with immunoprecipitation wash buffer (GTEN extraction buffer with 0.015% [v/v] Triton X-100 [Sigma-Aldrich]) and resuspended in 20 μl SDS loading dye. Proteins were eluted from the beads by heating for 10 min at 50°C for GFP or 95°C for FLAG or HA). The immunoprecipitates and input proteins were separated by SDS-PAGE and transferred onto polyvinylidene difluoride membranes using the Trans-Blot Turbo Transfer System (Bio-Rad, Munich), according to the manufacturer’s instructions. Blots were blocked with 5% skim milk in tris-buffered saline containing Tween20 for a minimum of 1 hr at room temperature. The immunoprecipitated proteins and input proteins were analyzed by immunoblotting with specific antibodies. Protein staining was imaged using the ImageQuant LAS 4000 luminescent imager (GE Healthcare Life Sciences, Piscataway, NJ).

### *In vitro* pull-down assay

Codon-optimized His:AVRblb2, CaM:GST, and CML36:GST were expressed in the BL21 (DE3) strain (RBC, Taiwan). The His:AVRblb2 protein was induced with 1 mM isopropyl β-D-1-thiogalactopyranoside (IPTG) at 37°C for 3 hr. The cells were centrifuged at 3,000 rpm for 15 min, resuspended in His-binding buffer (50 mM Tris-HCl [pH 8.0], 30 mM imidazole, 300 mM NaCl, 0.1 mM EDTA), and lysed by sonication. The lysate was centrifuged at 12,000 rpm for 10 min and filtered through a 0.45-μm syringe filter. The soluble proteins were purified using HisPur™ Cobalt Resin (Thermo Scientific, USA) and eluted in His-elution buffer (50 mM Tris-HCl [pH 8.0], 50 mM NaCl, 300 mM imidazole, and 0.1 mM EDTA). The purified protein was dialyzed in PBS buffer overnight. The CaM:GST and CML36:GST proteins were induced with 1 mM IPTG at 30°C overnight. The cells were centrifuged at 3,000 rpm for 15 min, resuspended in GST-binding buffer (50 mM Tris-HCl [pH 8.0], 150 mM NaCl, 0.1 mM EDTA), and lysed by sonication. The soluble proteins were purified using Glutathione Agarose (Thermo Scientific, USA) and eluted in GST-binding buffer containing 10 mM reduced glutathione. For the pull-down assay, His:AVRblb2 (2.5 μg) and GST-tagged prey protein (CaM or CML36, 2.5 μg each) were incubated in equilibrium buffer (20 mM Tris-HCl, [pH 7.5], 250 mM NaCl) containing 2 mM CaCl_2_ or 1 mM EDTA for 2.5 hr at 4°C with agitation. The bait protein alone served as a negative control. The incubated bait and prey proteins were added to a Ni-NTA agarose bead-containing column and incubated for 10 min on ice. The column was washed four times with equilibrium buffer containing 20 mM imidazole. The pull-down protein was eluted with elution buffer. Different concentrations of calcium were added to study the dependence of the pull-down affinity on calcium concentration.

### Quantitative polymerase chain reaction (PCR)

Total RNA was extracted from *N. benthamiana* leaves using TRIzol reagent (MRC, USA), and cDNA was generated using SuperScript III reverse transcriptase (Invitrogen, USA) and oligo (dT) primer following RNase-free DNase I treatment. Quantitative RT-PCR assays were conducted using PowerUP SYBR green master mix (Thermo, USA), according to the manufacturer’s instructions. PCR was performed using the primers listed in Table S1. The *N. benthamiana* EF1α gene was used as the internal reference gene for calculating the relative gene expression levels, which were calculated using the 2^-ΔΔCt^ algorithm (Rao et al., 2013).

### ROS measurements

ROS measurements were performed as previously described [46]. Four-week-old *N. benthamiana* plants for each genotype were excised into leaf discs (5-mm diameter) and floated 16 hr in 200 μl water in a 96-well plate. Leaf discs were soaked in 50 mM luminol containing 10 mg/ml horseradish peroxidase and 1 μM flg22 or water control. Beginning immediately after adding the luminol solution, luminescence was measured over a 45-min period using a multi-label plate reader (Varioskan LUX, ThermoFisher, USA). Data were analyzed using company software and Microsoft Excel.

### Pathogen inoculation

*P. infestans* T30-4 was grown on rye sucrose agar medium at 16°C in the dark. Zoospores were collected from 10 to 14-day-old cultures by flooding with cold water and incubating at 4°C for 60–90 min. For detached leaves, *P. infestans* infection assays were performed on the abaxial surface using droplet inoculation of zoospore suspensions (10 μl, 50,000 spores/ml). The phenotype was monitored for 7 days. Lesions were photographed 5 day post-inoculation using the Cy5 and Cy3 channels of the Azure 400 (Azure Biosystems, USA), and autofluorescent areas were analyzed using ImageJ software. At least three independent experiments were performed for this assay.

### Y2H assay

The Y2H assay was performed using the Matchmaker Gold system (Takara Bio, USA). Individual pGADT7 constructs or pGADT7 empty vector were introduced into the yeast strain Y187, and the cloned pGBKT7 plasmids were introduced into yeast strain Y2HGold. The yeast colonies containing both pGADT7 and pGBKT7 were selected on a synthetic defined (SD) medium without leucine and tryptophan (SD-L-T), and the interactions were tested on SD medium without histidine, leucine, and tryptophan (SD-H-L-T) supplemented with 0.2% adenine. Plates were incubated for 3 to 6 days at 28°C and then imaged. Each experiment was repeated a minimum of three times, yielding similar results. Commercial yeast constructs were used as positive (pGBKT7-53/pGADT7-T) and negative (pGBKT7-Lam/pGADT7-T) controls (Clontech, USA).

### BiFC assay

The full-length coding sequences of CNGCs and AVRblb2 were amplified and inserted into the pVYNE-35S or pSCYCE-35S vector. Agrobacterium strain GV3101 was transformed with the BiFC constructs before infiltrating four-week-old *N. benthamiana* leaves. Two days after infiltration, the samples were visualized using the Leica SP8 X (USA) confocal microscope equipped with a 40× water immersion objective. Fluorescence intensity was visualized using excitation at 488 nm and emission between 455 and 480 nm. Images were captured in multichannel mode with brightfield and processed using LAS X microscope software.

### Split-luciferase complementation assay

The split-luciferase complementation assay was performed as previously described (Luker and Luker, 2011). Briefly, cell suspensions of *A. tumefaciens* GV3101 carrying each construct were mixed in induction buffer (10 mM MES pH 5.7, 10 mM MgCl2, and 150 μM acetosyringone) at an OD600 of 0.5. Bacteria were infiltrated into *N. benthamiana* leaves, and the plants were grown in growth chambers for 2 days. Whole leaves or leaf disks were incubated with 1 mM luciferin. Luminescence was recorded using a microplate reader (Varioskan LUX, ThermoFisher, USA), and luminescence was quantified from twelve replicates of each sample.

### Virus-induced gene silencing (VIGS)

VIGS was performed in *N. benthamiana* as previously described (Chung et al., 2004). Suspensions of *A. tumefaciens* strain GV3101 carrying TRV-RNA1 and TRV-RNA2 containing the corresponding fragments from NbCNGC18, CaM, or CML36 were mixed in a 2:1 ratio in infiltration buffer (10 mM 2-[N-morpholine]ethanesulfonic acid [MES]; 10 mM MgCl2; and 150 μM acetosyringone, pH 5.6) to a final OD600 of 0.25. Two-week-old *N. benthamiana* plants were infiltrated with *A. tumefaciens*, and the upper leaves were used two to three weeks later for further assays.

## Supplemental figure legends

**Figure S1. Phylogenetic tree of NbCNGC proteins**.

The phylogenetic tree was created with the ClustalX program using the maxiAmino maximum likelihood method with 1000 bootstrap replications in MEGA7 (Tamura et al., 2011). NbCNGC18 is shown in blue.

**Figure S2. AVRblb2 paralogs suppress flg22-induced Ca2+ influx**.

(A) Co-IP of C-terminally GFP-tagged NbCNGC18-CT with N-terminally FLAG-tagged AVRblb2 homologs (PITG_20300, 20301, 20303, 18683, and 04090). Proteins were obtained by co-IP with GFP beads (GFP-IP). The experiments were performed more than three times using different pull-down conditions with similar results. (B) Time-series images of YC3.6-expressing *N. benthamiana* transgenic leaves after flg22 treatment. All AVRblb2 homolog-expressing leaves exhibited a similar, lower peak elevation in Ca^2+^ (ΔR/R0) compared to the GST control.

**Figure S3. Diverse PAMPs elicit rapid Ca2+ signals in *N. benthamiana***.

Calcium elevation is reduced after treatment with diverse PAMPs. Time-series of the response to 1 μM flg22, 1 μM elf18, or 100 ug/ml chitin in 3-week-old YC3.6-expressing *N. benthamiana* transgenic leaves.

**Figure S4. AVRblb2 target proteins are involved in flg22-induced Ca2+ influx**.

YC3.6 transgenic plants infiltrated with Agrobacterium carrying TRV2-GFP served as control plants. Slight dwarfism and leaf chlorosis were observed in CaM- and NbCNGC18-silenced plants. Silencing of NbCNGC18 or CML36 had low levels of flg22-induced Ca^2+^ influx, whereas flg22-induced Ca^2+^ influx was upregulated in CaM-silenced plants.

**Figure S5. AVRblb2 does not affect NbCNGC18 phosphorylation**.

(A-C) Co-IP was performed with agarose beads conjugated to GFP antibody (GFP-IP). Total protein extracts (input) and proteins obtained by co-IP were immunoblotted with antibodies indicated on the right. Phosphorylation was analyzed using anti-p-threonine (α-Thr) and anti-p-serine (α-Ser) antibodies. Similar results were observed in three biological replicates.

**Figure S6. AVRblb2 does not affect NbCNGC18 phosphorylation levels**.

(A) *In planta* BiFC assays showing the interactions of AVRblb2 fused to YFPn with NbCNGCs fused to SCFPc. NbCNGC3, NbCNGC4, NbCNGC18, NbCNGC19, and NbCNGC20 interacted with AVRblb2, producing yellow fluorescence. (B) NbCNGC18 interacts with other CNGCs *in planta*. BiFC analyses were carried out using 4-week-old *N. benthamiana* leaves. Confocal images were taken after 2 dpi. Scale bar = 20 μm.

**Figure S7. Expression of multiple NbCNGCs induced a defense response**.

Differential colonization of *P. infestans* T30-4 on 3-week-old *N. benthamiana* leaves. Leaves expressing NbCNGC18, NbCNGC20, and NbCNGC25 had more enhanced resistance against *P. infestans* compared to leaves expressing one or two CNGCs. Pictures were taken at 6 dpi.

**Figure S8. Proposed model for the role of AVRblb2 and NbCNGC18 heteromers in the calcium-based PTI signaling pathway in *N. benthamiana***.

AVRblb2 targets a subset of NbCNGCs (CNGC2, CNGC18, CNGC19, and CNGC20) and forms AVRblb2-CaM/CML36-CNGC complexes and suppresses flg22-induced Ca^2+^ influx by blocking the dissociation between CaM/CML36 and NbCNGC18. After PAMP recognition, several different CNGCs interact with NbCNGC18 to form a heterotetrameric complex modulated by AVRblb2.

## References

Boller, T., and Felix, G. (2009). A renaissance of elicitors: perception of microbe-associated molecular patterns and danger signals by pattern-recognition receptors. Annu Rev Plant Biol 60, 379–406. 10.1146/annurev.arplant.57.032905.105346.

Bozkurt, T.O., Schornack, S., Win, J., Shindo, T., Ilyas, M., Oliva, R., Cano, L.M., Jones, A.M., Huitema, E., van der Hoorn, R.A., and Kamoun, S. (2011). Phytophthora infestans effector AVRblb2 prevents secretion of a plant immune protease at the haustorial interface. Proc Natl Acad Sci U S A 108, 20832–20837. 10.1073/pnas.1112708109.

Brost, C., Studtrucker, T., Reimann, R., Denninger, P., Czekalla, J., Krebs, M., Fabry, B., Schumacher, K., Grossmann, G., and Dietrich, P. (2019). Multiple cyclic nucleotide-gated channels coordinate calcium oscillations and polar growth of root hairs. Plant J 99, 910–923. 10.1111/tpj.14371.

Chin, K., DeFalco, T.A., Moeder, W., and Yoshioka, K. (2013). The Arabidopsis cyclic nucleotide-gated ion channels AtCNGC2 and AtCNGC4 work in the same signaling pathway to regulate pathogen defense and floral transition. Plant Physiol 163, 611–624. 10.1104/pp.113.225680.

Chung, E., Seong, E., Kim, Y.C., Chung, E.J., Oh, S.K., Lee, S., and Choi, D. (2004). A method of high frequency virus-induced gene silencing in chili pepper. Molecules and Cells 17, 377–380.

Cui, H., Tsuda, K., and Parker, J.E. (2015). Effector-triggered immunity: from pathogen perception to robust defense. Annu Rev Plant Biol 66, 487–511. 10.1146/annurev-arplant-050213-040012.

Curran, A., Chang, I.F., Chang, C.L., Garg, S., Miguel, R.M., Barron, Y.D., Li, Y., Romanowsky, S., Cushman, J.C., Gribskov, M., et al. (2011). Calcium-dependent protein kinases from Arabidopsis show substrate specificity differences in an analysis of 103 substrates. Front Plant Sci 2, 36. 10.3389/fpls.2011.00036.

DeFalco, T.A., Marshall, C.B., Munro, K., Kang, H.G., Moeder, W., Ikura, M., Snedden, W.A., and Yoshioka, K. (2016). Multiple calmodulin-binding sites positively and negatively regulate Arabidopsis CYCLIC NUCLEOTIDE-GATED CHANNEL12. Plant Cell 28, 1738–1751. 10.1105/tpc.15.00870.

Dietrich, P., Moeder, W., and Yoshioka, K. (2020). Plant cyclic nucleotide-gated channels: New insights on their functions and regulation. Plant Physiol 184, 27–38. 10.1104/pp.20.00425.

Du, Y., Chen, X., Guo, Y., Zhang, X., Zhang, H., Li, F., Huang, G., Meng, Y., and Shan, W. (2021). Phytophthora infestans RXLR effector PITG20303 targets a potato MKK1 protein to suppress plant immunity. New Phytol 229, 501–515. 10.1111/nph.16861.

Fischer, C., DeFalco, T.A., Karia, P., Snedden, W.A., Moeder, W., Yoshioka, K., and Dietrich, P. (2017). Calmodulin as a Ca2+-sensing subunit of Arabidopsis cyclic nucleotide-gated channel complexes. Plant Cell Physiol 58, 1208–1221. 10.1093/pcp/pcx052.

Grant, M., Brown, I., Adams, S., Knight, M., Ainslie, A., and Mansfield, J. (2000). The RPM1 plant disease resistance gene facilitates a rapid and sustained increase in cytosolic calcium that is necessary for the oxidative burst and hypersensitive cell death. Plant J 23, 441–450. 10.1046/j.1365-313x.2000.00804.x.

Guo, M., Kim, P., Li, G., Elowsky, C.G., and Alfano, J.R. (2016). A bacterial effector co-opts calmodulin to target the plant microtubule network. Cell Host Microbe 19, 67–78. 10.1016/j.chom.2015.12.007.

Heese, A., Hann, D.R., Gimenez-Ibanez, S. J. M. A., He, K., Li, J., and Rathjen, J.P. (2007). The receptor-like kinase SERK3/BAK1 is a central regulator of innate immunity in plants. Proc. Natl. Acad. Sci. USA 104, 12217–12222.

Jones, J.D., and Dangl, J.L. (2006). The plant immune system. Nature 444, 323–329. 10.1038/nature05286.

Kadota, Y., Liebrand, T.W.H., Goto, Y., Sklenar, J., Derbyshire, P., Menke, F.L.H., Torres, M.A., Molina, A., Zipfel, C., Coaker, G., and Shirasu, K. (2019). Quantitative phosphoproteomic analysis reveals common regulatory mechanisms between effector- and PAMP-triggered immunity in plants. New Phytol 221, 2160–2175. 10.1111/nph.15523.

Ladwig, F., Dahlke, R.I., Stuhrwohldt, N., Hartmann, J., Harter, K., and Sauter, M. (2015). Phytosulfokine regulates growth in Arabidopsis through a response module at the plasma membrane that includes CYCLIC NUCLEOTIDE-GATED CHANNEL17, H+-ATPase, and BAK1. Plant Cell 27, 1718–1729. 10.1105/tpc.15.00306.

Luker, G.D., and Luker, K.E. (2011). Luciferase protein complementation assays for bioluminescence imaging of cells and mice. Methods Mol Biol 680, 29–43. 10.1007/978-1-60761-901-7_2.

Moeder, W., Phan, V., and Yoshioka, K. (2019). Ca2+ to the rescue - Ca2+ channels and signaling in plant immunity. Plant Sci 279, 19–26. 10.1016/j.plantsci.2018.04.012.

Naveed, Z.A., Bibi, S., and Ali, G.S. (2019). The Phytophthora RXLR effector Avrblb2 modulates plant immunity by interfering with Ca2+ signaling pathway. Front Plant Sci 10, 374. 10.3389/fpls.2019.00374.

Oh, S.K., Young, C., Lee, M., Oliva, R., Bozkurt, T.O., Cano, L.M., Win, J., Bos, J.I., Liu, H.Y., van Damme, M., et al. (2009). In planta expression screens of Phytophthora infestans RXLR effectors reveal diverse phenotypes, including activation of the Solanum bulbocastanum disease resistance protein Rpi-blb2. Plant Cell 21, 2928–2947. 10.1105/tpc.109.068247.

Oliva, R.F., Cano, L.M., Raffaele, S., Win, J., Bozkurt, T.O., Belhaj, K., Oh, S.K., Thines, M., and Kamoun, S. (2015). A recent expansion of the RXLR effector gene Avrblb2 is maintained in global populations of Phytophthora infestans indicating different contributions to virulence. Mol Plant Microbe Interact 28, 901–912. 10.1094/MPMI-12-14-0393-R.

Pan, Y., Chai, X., Gao, Q., Zhou, L., Zhang, S., Li, L., and Luan, S. (2019). Dynamic interactions of plant CNGC subunits and calmodulins drive oscillatory Ca2+ channel activities. Dev Cell 48, 710–725 e715. 10.1016/j.devcel.2018.12.025.

Rao, X., Huang, X., Zhou, Z., and Lin, X. (2013). An improvement of the 2^ (–delta delta CT) method for quantitative real-time polymerase chain reaction data analysis. Biostatistics, Bioinformatics and Biomathematics 3, 71.

Saand, M.A., Xu, Y.P., Li, W., Wang, J.P., and Cai, X.Z. (2015). Cyclic nucleotide gated channel gene family in tomato: genome-wide identification and functional analyses in disease resistance. Front Plant Sci 6, 303. 10.3389/fpls.2015.00303.

Tamura, K., Peterson, D., Peterson, N., Stecher, G., Nei, M., and Kumar, S. (2011). MEGA5: molecular evolutionary genetics analysis using maximum likelihood, evolutionary distance, and maximum parsimony methods. Mol Biol Evol 28, 2731–2739. 10.1093/molbev/msr121.

Tian, W., Hou, C., Ren, Z., Wang, C., Zhao, F., Dahlbeck, D., Hu, S., Zhang, L., Niu, Q., Li, L., et al. (2019). A calmodulin-gated calcium channel links pathogen patterns to plant immunity. Nature 572, 131–135. 10.1038/s41586-019-1413-y.

Tian, W., Wang, C., Gao, Q., Li, L., and Luan, S. (2020). Calcium spikes, waves and oscillations in plant development and biotic interactions. Nat Plants 6, 750–759. 10.1038/s41477-020-0667-6.

Wang, J., Liu, X., Zhang, A., Ren, Y., Wu, F., Wang, G., Xu, Y., Lei, C., Zhu, S., Pan, T., et al. (2019). A cyclic nucleotide-gated channel mediates cytoplasmic calcium elevation and disease resistance in rice. Cell Res 29, 820–831. 10.1038/s41422-019-0219-7.

Whisson, S.C., Boevink, P.C., Moleleki, L., Avrova, A.O., Morales, J.G., Gilroy, E.M., Armstrong, M.R., Grouffaud, S., van West, P., Chapman, S., et al. (2007). A translocation signal for delivery of oomycete effector proteins into host plant cells. Nature 450, 115–118. 10.1038/nature06203.

Xu, G., Moeder, W., Yoshioka, K., and Shan, L. (2022). A tale of many families: Calcium channels in plant immunity. Plant Cell. 10.1093/plcell/koac033.

Yoshioka, K., Moeder, W., Kang, H.G., Kachroo, P., Masmoudi, K., Berkowitz, G., and Klessig, D.F. (2006). The chimeric Arabidopsis CYCLIC NUCLEOTIDE-GATED ION CHANNEL11/12 activates multiple pathogen resistance responses. Plant Cell 18, 747–763. 10.1105/tpc.105.038786.

Yu, X., Xu, G., Li, B., de Souza Vespoli, L., Liu, H., Moeder, W., Chen, S., de Oliveira, M.V.V., Ariadina de Souza, S., Shao, W., et al. (2019). The receptor kinases BAK1/SERK4 regulate Ca2+ channel-mediated cellular homeostasis for cell death containment. Curr Biol 29, 3778–3790 e3778. 10.1016/j.cub.2019.09.018.

Yuan, M., Jiang, Z., Bi, G., Nomura, K., Liu, M., Wang, Y., Cai, B., Zhou, J.M., He, S.Y., and Xin, X.F. (2021). Pattern-recognition receptors are required for NLR-mediated plant immunity. Nature 592, 105–109. 10.1038/s41586-021-03316-6.

Zhao, C., Tang, Y., Wang, J., Zeng, Y., Sun, H., Zheng, Z., Su, R., Schneeberger, K., Parker, J.E., and Cui, H. (2021). A mis-regulated cyclic nucleotide-gated channel mediates cytosolic calcium elevation and activates immunity in Arabidopsis. New Phytol 230, 1078–1094. 10.1111/nph.17218.

Zhao, Y., Liu, W., Xu, Y.P., Cao, J.Y., and Braam, J.C. X. Z. (2013). Genome-wide identification and functional analyses of calmodulin genes in Solanaceous species. BMC Plant Biol 13, 1–15.

Zheng, X., McLellan, H., Fraiture, M., Liu, X., Boevink, P.C., Gilroy, E.M., Chen, Y., Kandel, K., Sessa, G., Birch, P.R., and Brunner, F. (2014). Functionally redundant RXLR effectors from Phytophthora infestans act at different steps to suppress early flg22-triggered immunity. PLoS Pathog 10, e1004057. 10.1371/journal.ppat.1004057.

Zheng, X., Wagener, N., McLellan, H., Boevink, P.C., Hua, C., Birch, P.R.J., and Brunner, F. (2018). Phytophthora infestans RXLR effector SFI5 requires association with calmodulin for PTI/MTI suppressing activity. New Phytol 219, 1433–1446. 10.1111/nph.15250.

Zhou, L., Lan, W., Jiang, Y., Fang, W., and Luan, S. (2014). A calcium-dependent protein kinase interacts with and activates a calcium channel to regulate pollen tube growth. Mol Plant 7, 369–376. 10.1093/mp/sst125.

