## Supplemental figures for "Oomycete effector AVRblb2 targets cyclic nucleotide-gated channels through calcium sensors to suppress pattern-triggered immunity"

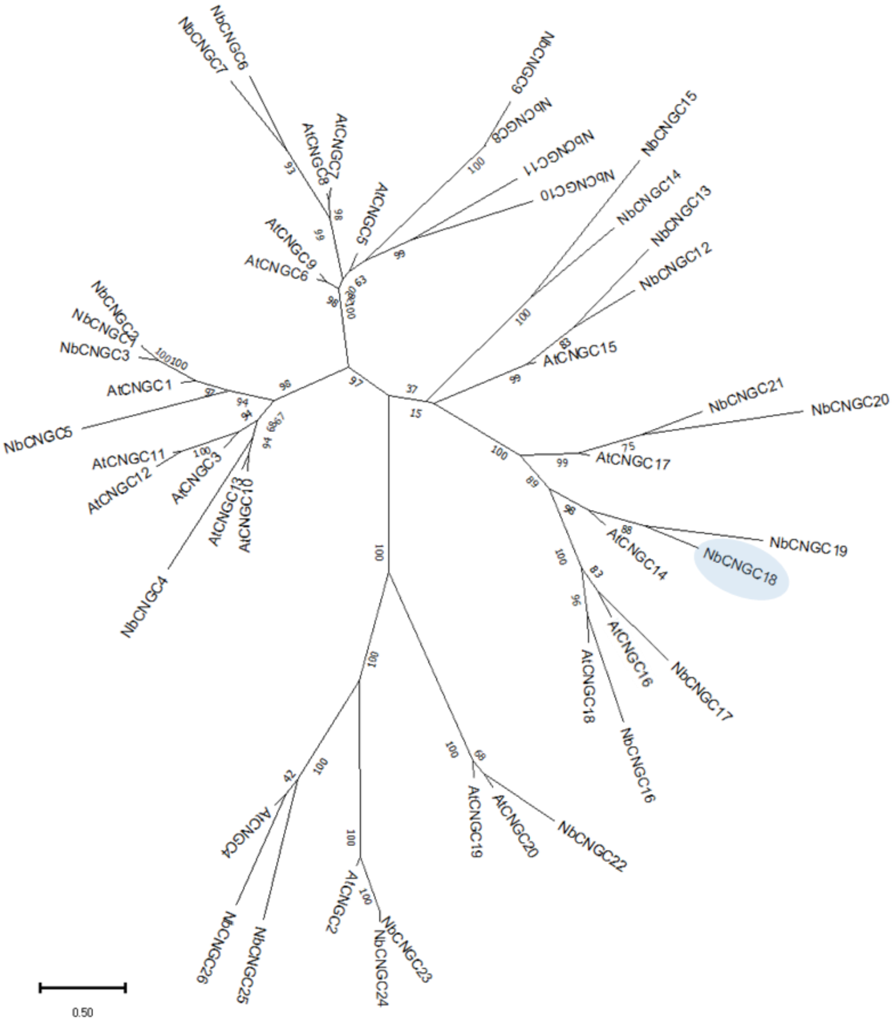


**Figure S1. Phylogenetic tree of the NbCNGC proteins.** The tree was created using Clustalx program by maxiAmino Maximum Likelihood (ML) method with 1000 bootstrap replication in MEGA7 (Tamura and Nei, 1993; Tamura et al., 2011). NbCNGC18/19 were shown in blue.

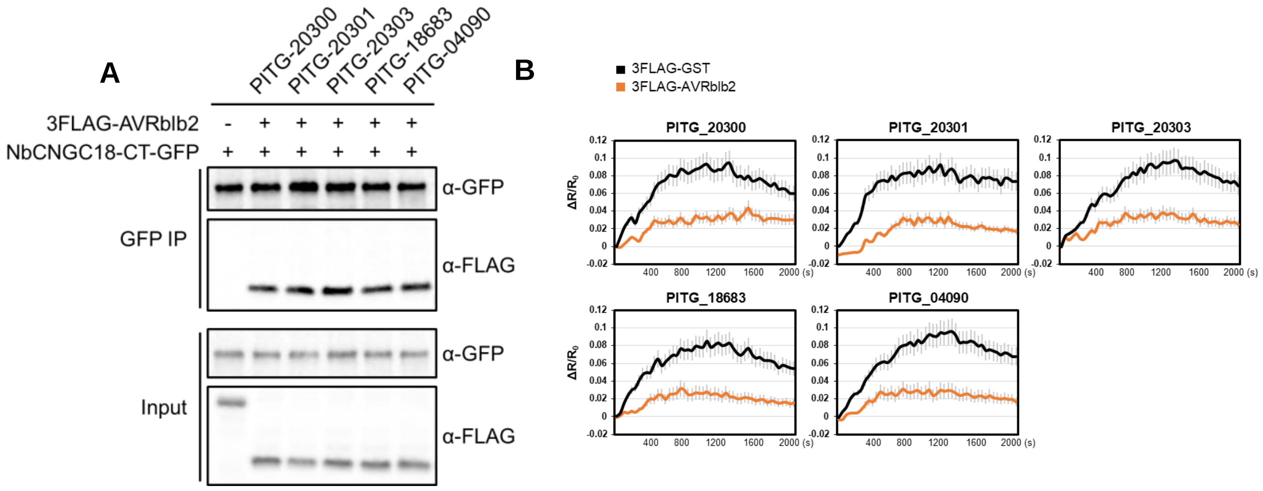


**Figure S2. AVRblb2 homologs suppress flg22-induced $Ca^{2+}$ influx.** (A) Co-IP assay of C-terminally GFP tagged NbCNGC18-CT with N-terminally FLAG tagged AVRblb2 homologs (PITG_20300, 20301, 20303, 18683, 04090). Proteins obtained by co-IP with GFP beads (GFP IP) and the experiment was performed more than three times under different pulldown conditions with similar results. (B) Times-series images of YC3.6 *N. benthamiana* transgenic leaves after flg22 treatment. Note all AVRblb2 homolog expressed leaves exhibit a similar, lower peak elevation in $Ca^{2+}$ ($\Delta R/R0$) compared to GST control.

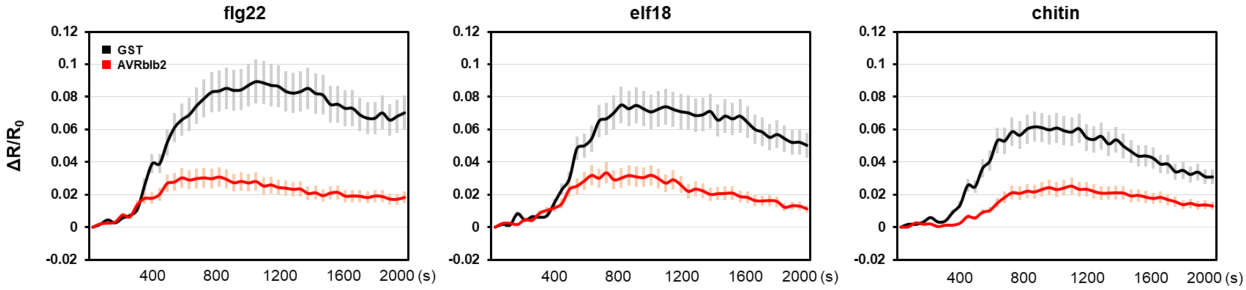


**Figure S3. Diverse PAMPs elicit rapid $Ca^{2+}$ signals in *N. benthamiana*.** Calcium elevation is reduced after treatment with diverse PAMPs. Time-series of response in 3 week-old YC3.6 *N. benthamiana* transgenic leaves treated with 1 μM flg22, 1 μM elf18 and 100 ug/ml chitin respectively.

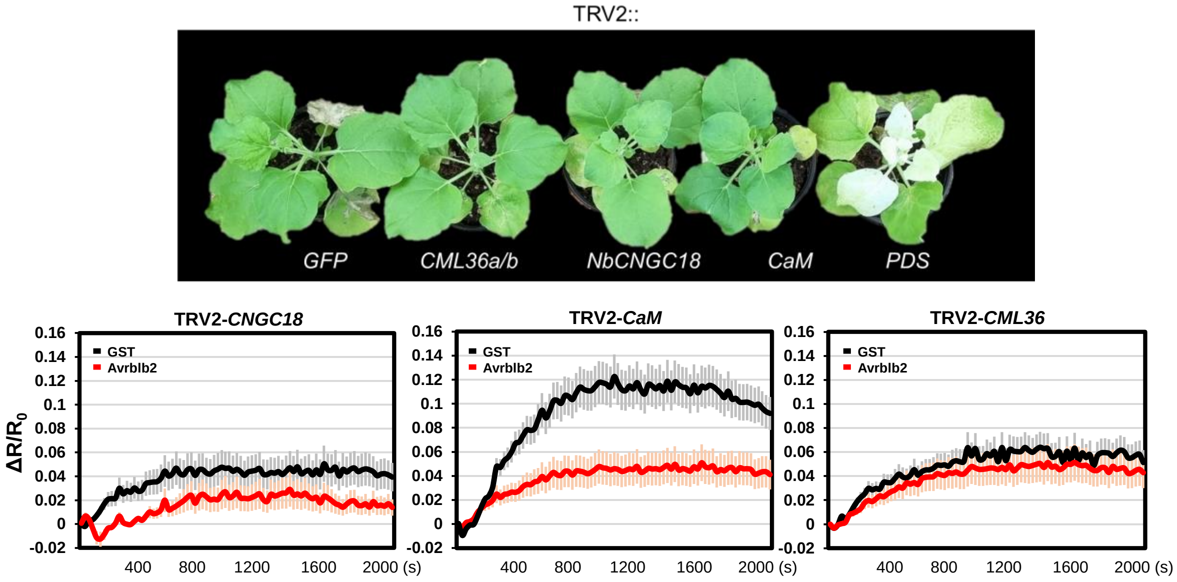


**Figure S4. AVRblb2 target proteins are involved in flg22-induced $Ca^{2+}$ influx.** YC3.6 transgenic plant infiltrated with Agrobacterium carring an TRV2-GFP control served as control plants. Slight dwarfism and leaf chlorosis were observed in CaM and NbCNGC18 silenced plant. Silencing of NbCNGC18 or CML36 plant showed a low level of flg22-induced $Ca^{2+}$ influx, but up-regulated in CaM silencing plant.

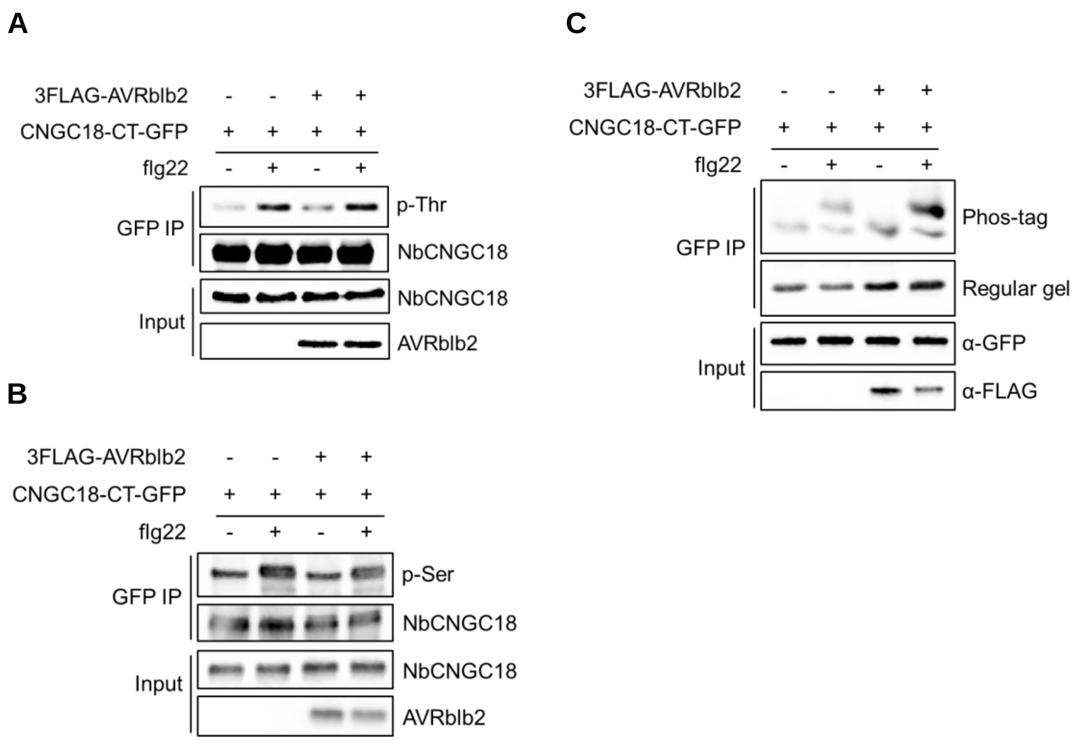


**Figure S5. AVRblb2 does not affect the phosphorylation level of NbCNGC18. (**A-C) Co-IP was performed with agarose beads conjugated to GFP (GFP IP) antibodies. Total protein extracts (Input) and proteins obtained by co-IP were immunoblotted with appropriate antibody labelled on the right. The tyrosine phosphorylation was probed by anti-pThreonine (α-Thr) and anti-Serine (α-Ser) antibody. Similar results were observed in three biological replicates.

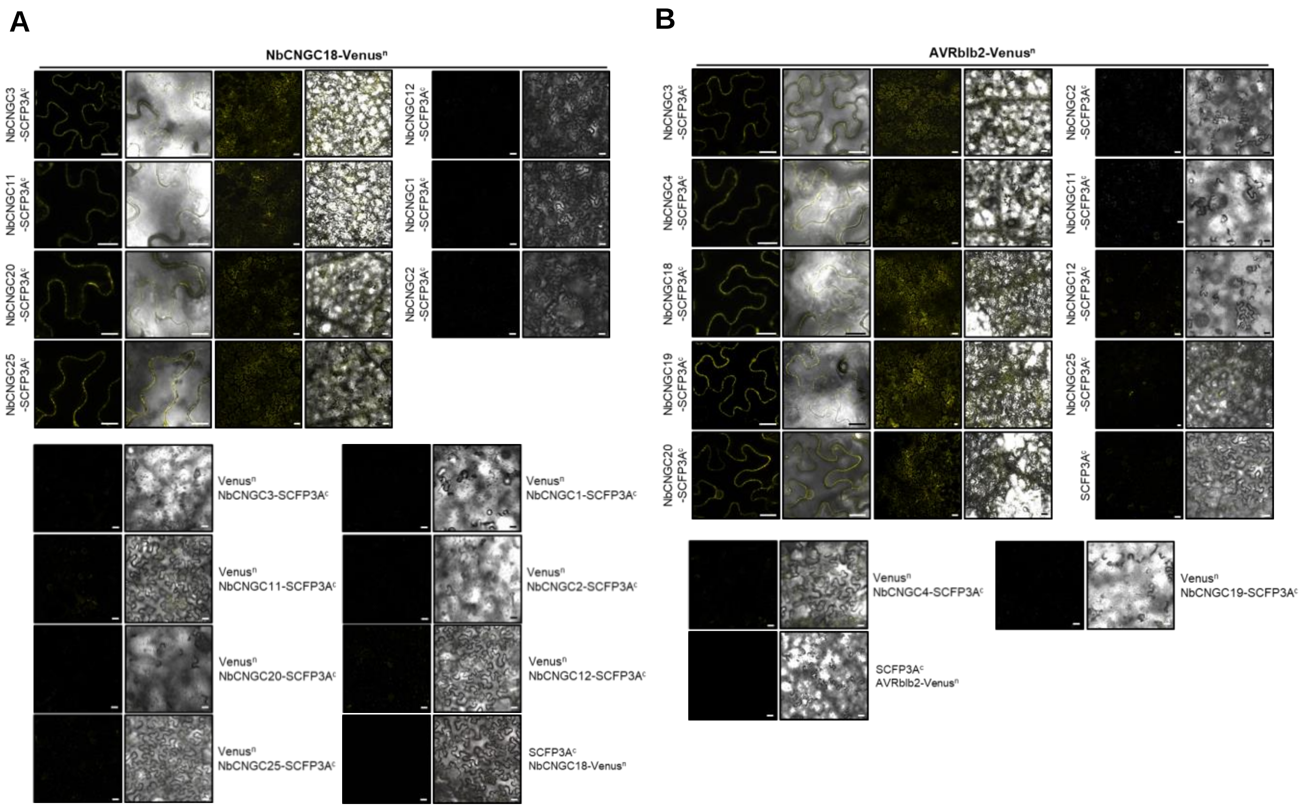


**Figure S6. AVRblb2 does not affect the phosphorylation level of NbCNGC18.** (A) *In planta* BiFC assays showing interaction of AVRblb2 fused to YFPn with NbCNGCs fused to SCFPc. NbCNGC3/4/18/19/20 show interaction with AVRblb2 and give yellow fluorescence. (B) NbCNGC18 interact with other CNGCs *in planta*. BiFC analyses were carried out using 4 weeks-old *N. benthamiana* leaves. Scale bar represents 20 µm, and the confocal images were taken after 2 dpi.

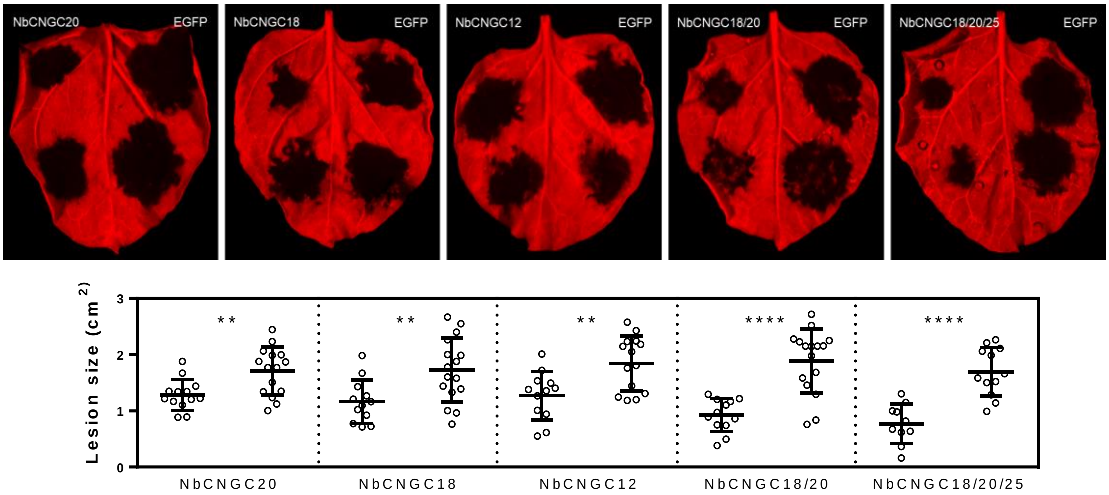


**Figure S7. Multiple expression of NbCNGCs could induce defense response.** Differential colonization of P. infestans T30-4 on N. benthamiana leaves (3 wk old). NbCNGC18/20/25 expressed leaves showed a more enhanced resistance against P. infestans compared to single or double expression of CNGC. Pictures were taken at 6 dpi.

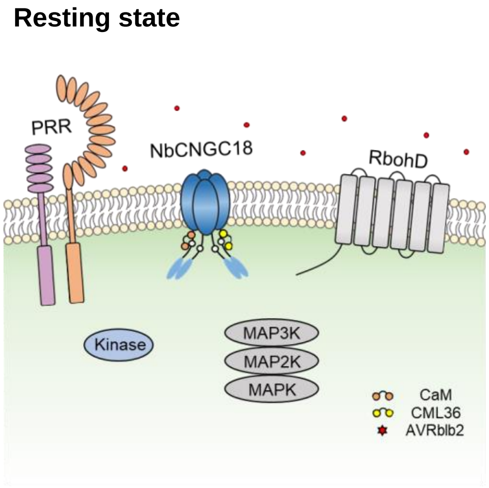


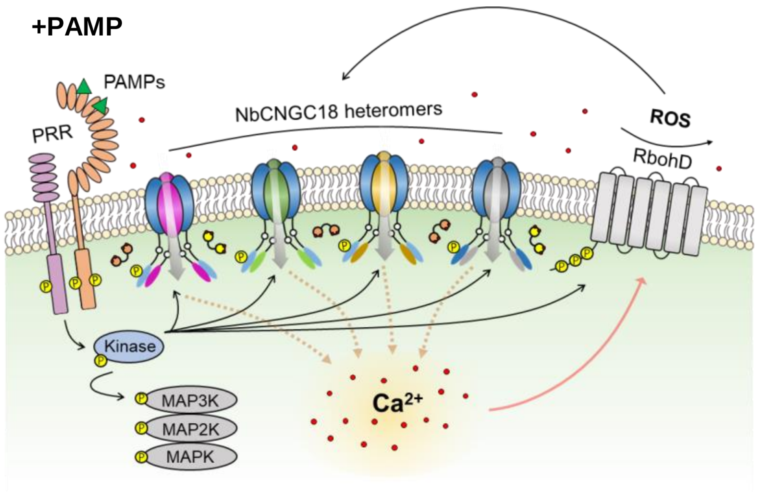


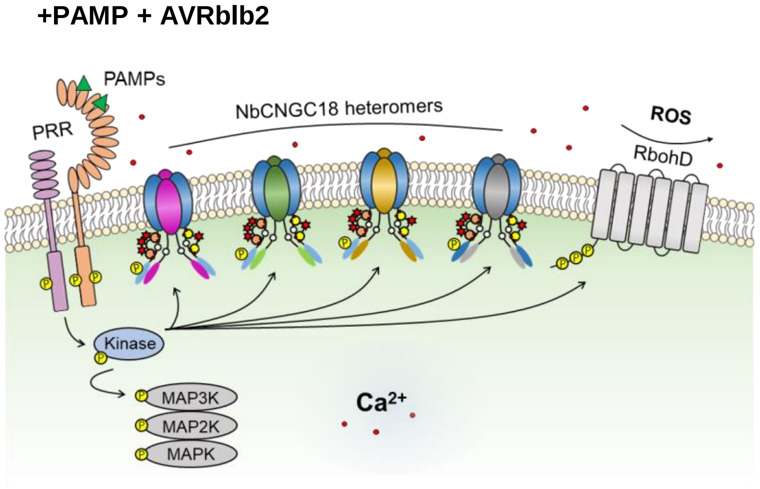


**Figure S8. Proposed model for the role of AVRblb2 and NbCNGC18 heteromers in the calcium based PTI signaling pathway in *N. benthamiana*.** AVRblb2 targets a subset of NbCNGCs (CNGC2, CNGC18, CNGC19, and CNGC20) and forms AVRblb2-CaM/CML36-CNGC complexes and suppresses flg22-induced $Ca^{2+}$ influx by blocking the dissociation between CaM/CML36 and NbCNGC18. After PAMP recognition, several different CNGCs interact with NbCNGC18 to form a heterotetrameric complex modulated by AVRblb2.
